# Shared neural resources of rhythm and syntax: An ALE Meta-Analysis

**DOI:** 10.1101/822676

**Authors:** Matthew Heard, Yune S. Lee

**Affiliations:** Department of Speech and Hearing Sciences, The Ohio State University; Neuroscience Graduate Program, The Ohio State University; Chronic Brain Injury, Discovery Theme, The Ohio State University, Columbus, OH 43210, USA

**Keywords:** rhythm, beat, meter, temporal processing, merge, movement, reanalysis, syntax, activation likelihood estimate, meta-analysis, fMRI

## Abstract

A growing body of evidence has highlighted behavioral connections between musical rhythm and linguistic syntax, suggesting that these may be mediated by common neural resources. Here, we performed a quantitative meta-analysis of neuroimaging studies using activation likelihood estimate (ALE) to localize the shared neural structures engaged in a representative set of musical rhythm (rhythm, beat, and meter) and linguistic syntax (merge movement, and reanalysis). Rhythm engaged a bilateral sensorimotor network throughout the brain consisting of the inferior frontal gyri, supplementary motor area, superior temporal gyri/temporoparietal junction, insula, the intraparietal lobule, and putamen. By contrast, syntax mostly recruited the left sensorimotor network including the inferior frontal gyrus, posterior superior temporal gyrus, premotor cortex, and supplementary motor area. Intersections between rhythm and syntax maps yielded overlapping regions in the left inferior frontal gyrus, left supplementary motor area, and bilateral insula—neural substrates involved in temporal hierarchy processing and predictive coding. Together, this is the first neuroimaging meta-analysis providing detailed anatomical overlap of sensorimotor regions recruited for musical rhythm and linguistic syntax.

## 1. Introduction

Both music and language tasks require efficient analysis of sequential structures in a given sequence. In the music domain, this operation is especially important for understanding the rhythm of a song. Rhythm refers to the temporal pattern of accented and unaccented auditory events present in music (Vuust and Witek, 2014). From the rhythm, listeners are able to extract the beat, the isochronous psychological event that drives the music forward (Grahn, 2012; Nguyen et al., 2018). Beat, also called pulse, in turn gives rise to meter–perceptual patterns of “strong” and “weak” beats—that can be internally or externally driven (**Figure 1A**; Iversen et al., 2009; Nozaradan et al., 2011). We consider these two derivatives as “rhythm” in the current meta-analysis. In the language domain, similar neural mechanisms may be at play in binding and moving around syntactic phrases in a given sentence (Kotz et al., 2009; Rothermich et al., 2012; Schmidt-Kassow and Kotz, 2008a). Notably, it has been suggested that natural grammar learning is based upon temporal context of spoken sentences. For instance, infants can use prosodic cues to identify syntactic boundaries (Fernald and McRoberts, 1996), and children adhere to strong metrical sequences while listening to sentences (Moritz et al., 2013; Strait et al., 2011). It has also been shown that metric patterns of speech facilitate comprehension of syntactically challenging sentences (Roncaglia-Denissen et al., 2013). Conversely, failure to detect timing cues may give rise to speech and language disorders such as dyslexia and specific language impairment (SLI; Goswami, 2011). Indeed, there are ample reports regarding deficits in rhythm and grammar in dyslexia and SLI, suggesting dysfunctional temporal processing is responsible for these developmental language disorders (Corriveau and Goswami, 2009; Gordon et al., 2015b; Goswami et al., 2013; Huss et al., 2011; Thomson and Goswami, 2008). Accordingly, rhythm training has been utilized as speech and language intervention programs for these populations (Bedoin et al., 2016; Bhide et al., 2013; Flaugnacco et al., 2015; Gordon et al., 2015b; Ozernov-Palchik et al., 2018; Przybylski et al., 2013).

**Figure 1:**
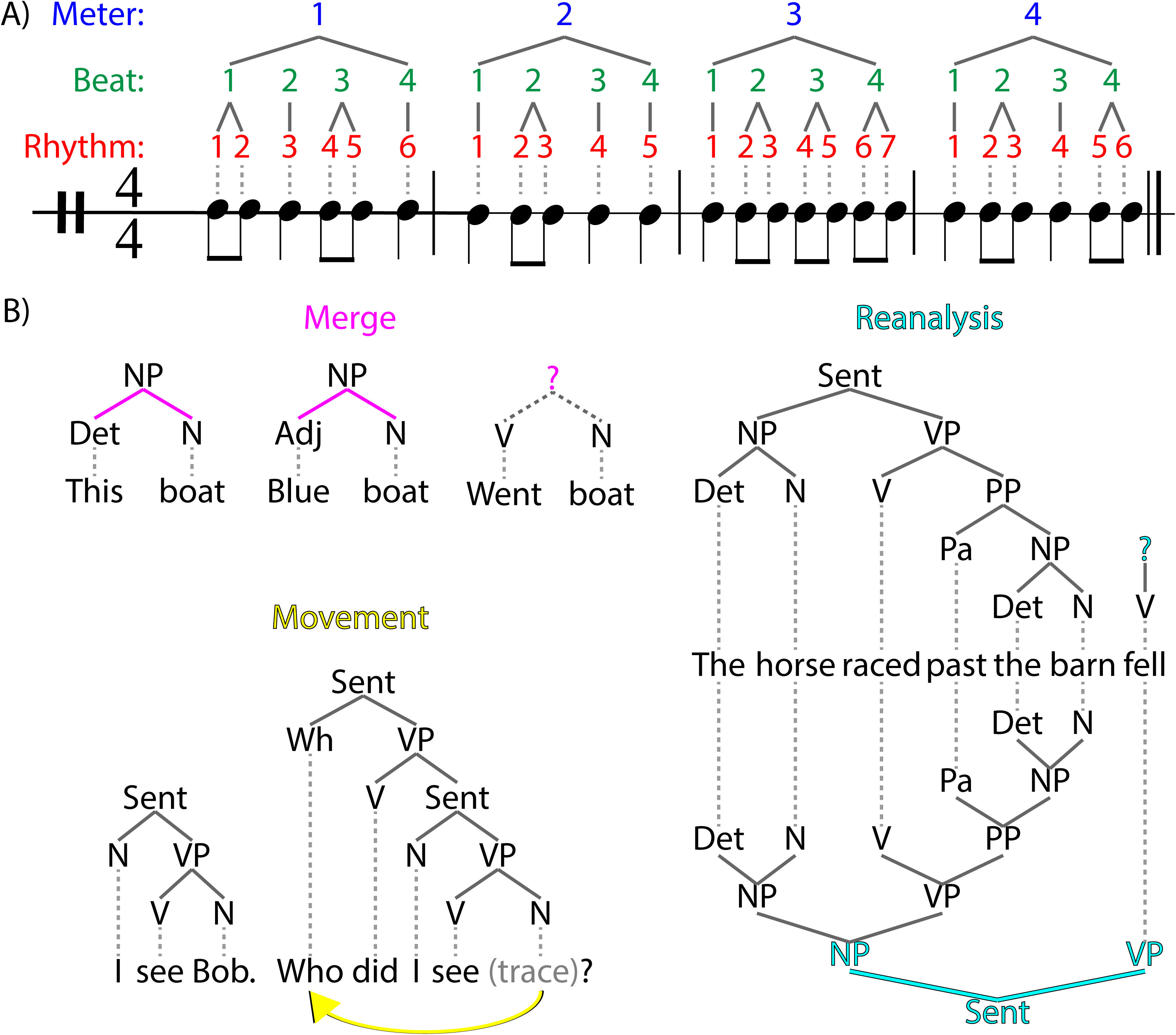
Schematics of rhythm and syntax. (A) An example music sequence consisting of quarter and eighth notes. Rhythms (in red) are the pattern of onsets perceived by the listener. Beat and meter (in green and blue) are extracted from the rhythms by the listener. (B) Three representative examples of syntax explored in the present meta-analysis. Merge brings together words or smaller phrases into larger phrases. Movement processes dependent nodes that are often found in wh-questions. Reanalysis occurs when extracting complicated grammatical roles resolving ambiguous word orders, such as in the garden path sentence exhibited here. NP: noun phrase; Sent: sentence; Det: determinant; N: noun; Adj: adjective; Wh: question word; VP: verb phrase; Pa: participle.

Within linguistic syntax, there are several sub-operations including merge, movement, reanalysis, syntactic surprisal (also known as prediction), morphosyntactic task, and others. In the present article, our scope is limited to merge, movement, and reanalysis (**Figure 1B**) as these were the only domains with enough experiments that passed our criteria of inclusion (see Methods 2.2 for more details). Merge involves combining words and phrases into larger syntactic building units (e.g. for + him = {for him}) (Chomsky, 1995; Zaccarella et al., 2017). Movement involves shifting phrases to fill dependent nodes such as traces in wh-questions (e.g. “What concert did you go to?”) (Grodzinsky and Santi, 2008; Santi and Grodzinsky, 2007a, 2007b). Reanalysis is the process of listeners revising previously constructed phrases, such as in garden path (e.g. “the horse raced passed the barn fell”) and object-relative (e.g. “boys that girls help are nice”) sentences (Caplan and Waters, 1999; Sturt and Crocker, 1996).

While both rhythm and syntax networks have been extensively studied by neuroimaging, the degree to which these networks overlap with or are segregated from each other remains to be determined. This may be a timely and important question given that there has been emerging evidence of behavioral connections between music and language (Gordon et al., 2015a). To this end, we performed a series of neuroimaging meta-analyses using activation likelihood estimate (ALE) (Chein et al., 2002; Eickhoff et al., 2012, 2009; Turkeltaub et al., 2012) on a set of experiments examining musical rhythm (rhythm, beat, meter) and linguistic syntax (merge, movement, reanalysis). By considering various types of rhythm and syntax, we avoided limiting the scope of our ALE findings to particular processes. ALE was developed independently by two groups (Chein et al., 2002; Turkeltaub et al., 2002) and has been widely used in the neuroimaging community to identify brain regions that are consistently implicated across numerous studies for particular sensory/cognitive processes. We first attempted to identify the core resources of rhythm and syntax independently by performing ALE analyses within each domain. To ensure that the activation maps computed equally represented each sub-process of rhythm (rhythm, meter, beat) and syntax (merge, movement, reanalysis), the number of experiments included from each sub-process was matched. Then, we identified overlapping regions between these core rhythm and syntax regions. Together, the present article affords a comprehensive picture of the common neural structures engaged in rhythm and syntax processing.

## 2. Methods

### 2.1 ALE Meta-analysis

In ALE, foci of activation are modeled as three-dimensional Gaussian distributions of probability in order to capture some of the geographical uncertainty that is inherent to fMRI and PET experiments:

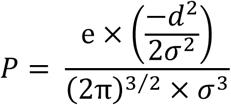

This equation transforms each focus into a three-dimensional Gaussian distribution. Every voxel in the brain is assigned a probability *P* based on *d*, the Euclidean distance between each voxel and the nearest coordinate drawn from each experiment, and *σ*, the degree of noise, which depends on the number of subjects from the experiment (Eickhoff et al., 2009; Wiener et al., 2010). By repeating this procedure across all reported foci from each experiment, modeled activation (MA) scores are computed (Turkeltaub et al., 2012). Then, the ALE score of each voxel is determined by summing all of the MA scores from all experiments. Finally, the resulting map is tested against the null distribution in which all foci are randomly and independently scattered through the gray matter of the brain (Eickhoff et al., 2012; Turkeltaub et al., 2012).

### 2.2 Literature Search

In order to find experiments to include in this meta-analysis, a literature search was performed on October 29^th^, 2018 using PubMed. The following searches were performed to locate relevant papers related to rhythm: “rhythm AND (fMRI OR “Functional Magnetic Resonance”) NOT cardiac NOT sleep”, “rhythm AND (PET OR “Positron Emission Tomography”)”, “meter AND (fMRI OR “Functional Magnetic Resonance”)”, “meter AND (PET OR “Positron Emission Tomography”)”, “beat AND (fMRI OR “Functional Magnetic Resonance”)”, “beat AND (PET OR “Positron Emission Tomography”)”. Similarly, the following searches were performed to locate papers related to syntax processing: “(fMRI OR functional magnetic resonance imaging OR PET OR positron emission tomography) AND grammatical”, “(fMRI OR functional magnetic resonance imaging OR PET OR positron emission tomography) AND syntactic”, “(fMRI OR functional magnetic resonance imaging OR PET OR positron emission tomography) AND grammar”, “(fMRI OR functional magnetic resonance imaging OR PET OR positron emission tomography) AND syntax”. Additional rhythm papers were located by reviewing the citations of these nine review papers: (Fitch, 2013; Geiser et al., 2014; Grahn, 2012; Kotz et al., 2018; Merchant et al., 2015; Pearce and Rohrmeier, 2012; Repp and Su, 2013; Teki, 2016; Teki et al., 2012). The full process of elimination is summarized according to PRISMA standards within Supplementary Materials S1 and S2 (Moher et al., 2009).

First, papers were screened based on titles. Due to the large quantity of papers identified by the PubMed database search, an in-house MATLAB script (2017a) was assembled to remove papers that were not in English or contained certain keywords in titles that are associated with EEG/MEG methods, clinical populations, developmental research, aging participants, or non-human subjects. The remaining papers were manually inspected to remove additional off-target results. Eligibility was assessed based on whether the paper reported foci of activation from whole-brain analyses and used fMRI acquisition paradigms that achieved whole brain coverage. Furthermore, experiments were excluded from analysis if they failed to control for domain-agnostic task-general cognitive resources such as working memory. Results contrasting musicians and non-musicians were also excluded. Since GingerALE software generates MA scores per each set of unique subjects in a study, papers reporting results from two independent populations count as two separate experiments (Chen et al., 2007; Goucha and Friederici, 2015), and multiple papers that report data from the same subjects (Vuust et al., 2011, 2006) are considered one experiment (Turkeltaub et al., 2012).

Initially, the rhythm literature search revealed a total of 32 experiments: 7 rhythm, 6 meter, 13 beat, and 6 studies that conflated rhythm, meter, and beat. Unfortunately, there were not enough papers within the scope of the present analysis to allow for running rhythm, meter, and beat as independent samples of experiments. To ensure that the rhythm analysis equally revealed areas associated with beat, meter, and rhythm processing, we down-sampled the number of experiments to match the number of studies from the meter category. Only the most recent experiments from the beat and rhythm categories were included. Accordingly, only the six newest experiments of each category were included for a total of 24 experiments engaging rhythm, beat, meter, and a mixture of these processes (**Table 1**). For the syntax analysis, the literature search discovered 50 syntax experiments: 16 experiments on merge, 13 experiments on movement, and 20 experiments on reanalysis (**Table 2**). For the overall syntax analysis, we included the 13 most recent papers from each category and ran them as a single population to ensure the activation revealed was associated with equal parts from merge, movement, and reanalysis processing. Because each subset had enough experiments, we were able to perform ALE analyses separately on merge, movement, and reanalysis experiments in addition to the general syntax domain. A full table listing the precise contrast and foci included from each experiment can be found in the Supplementary Materials Table 1.

**Table 1:**
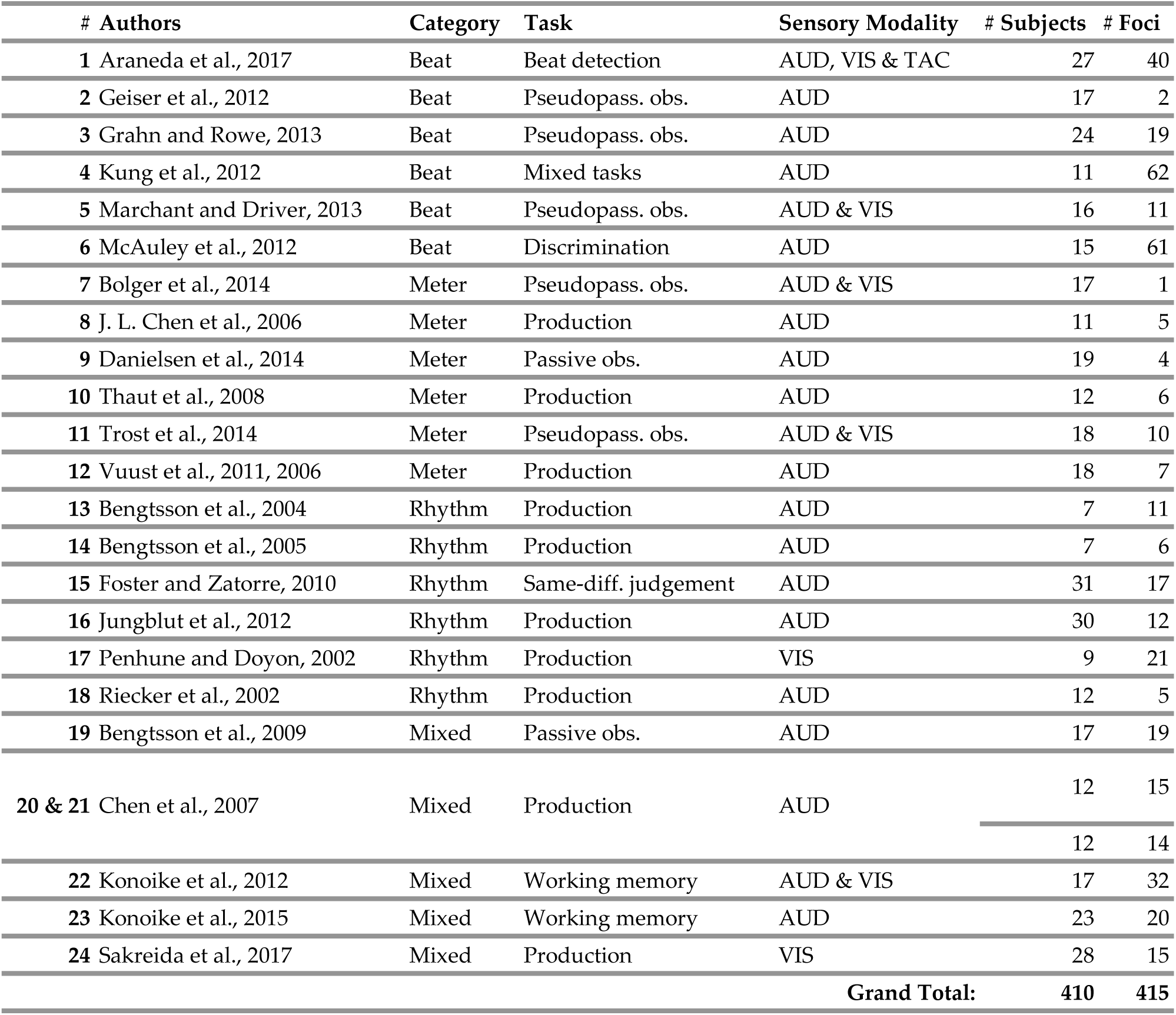
All experiments included in the rhythm ALE analysis. AUD: auditory; VIS: visual; TAC: tactile; Pseudopass. obs.: pseudopassive observation; Passive obs: passive observation; Same-diff. judgement: same-different judgement.

**Table 2:**
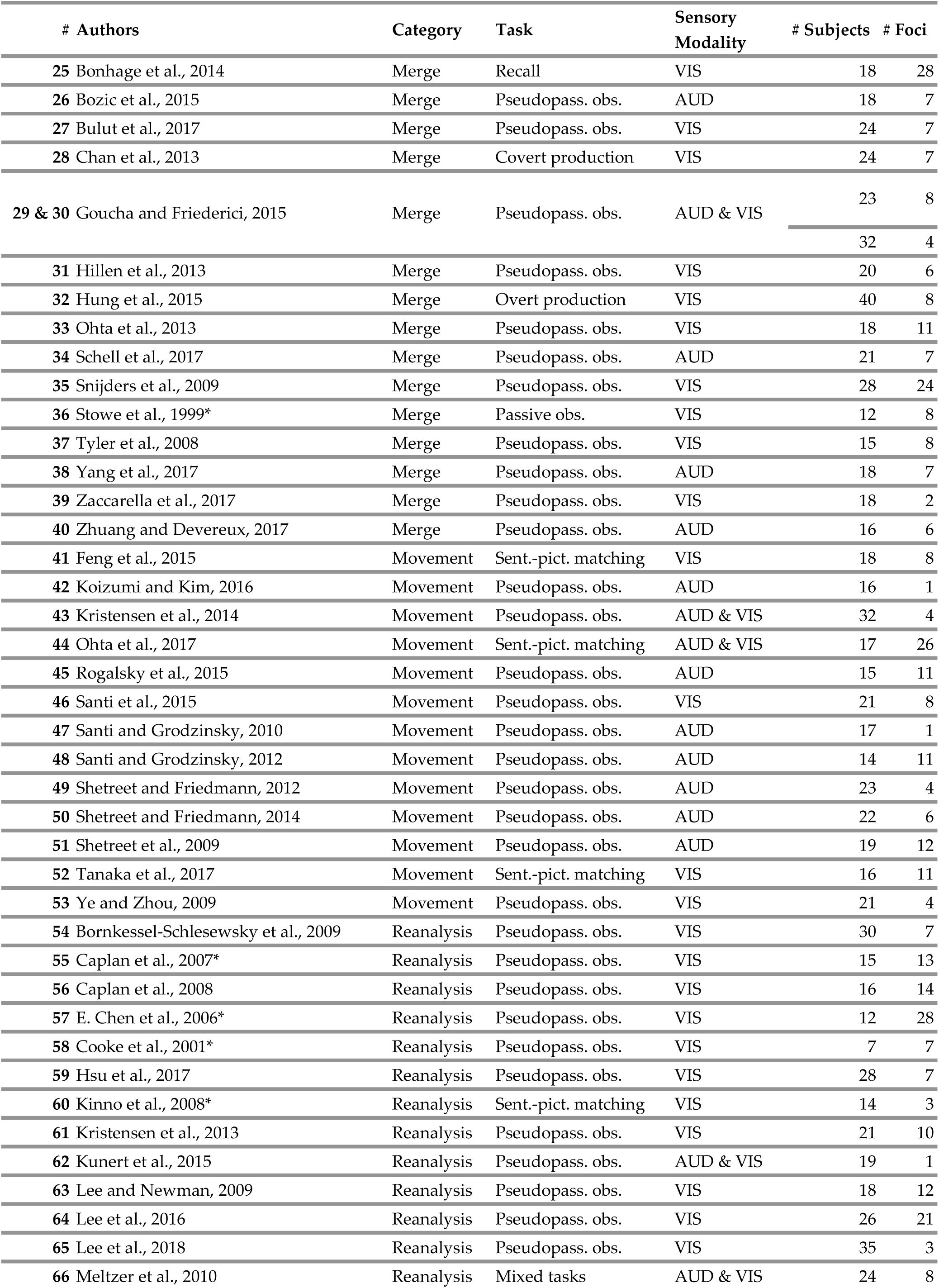
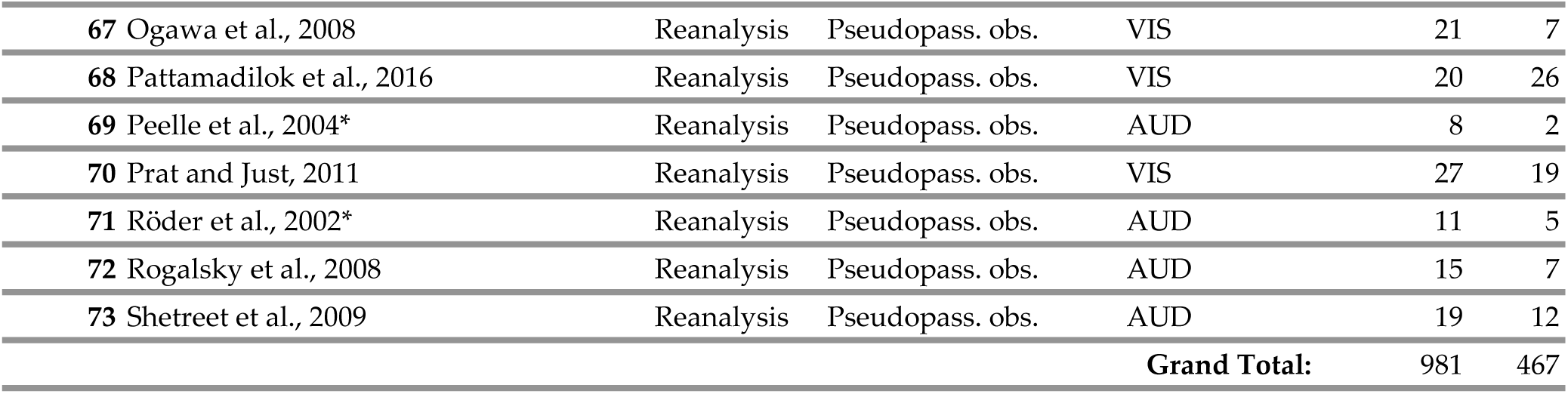
All experiments included in the syntax ALE analysis. VIS: visual; AUD: auditory; Pseudopass. obs.: pseudopassive observation; Passive obs.: passive observation; Sent.-pict. matching: sentence-picture matching. Asterisks indicate studies that were excluded when the syntax experiments were run as one sample.

### 2.3 Analysis Procedure

ALE analyses were performed on the rhythm and syntax foci sets independently using GingerALE (version 2.3.6, brainmap.org) (Eickhoff et al., 2012, 2009; Turkeltaub et al., 2012). To this end, foci originally reported in Talairach coordinates were transformed into common MNI space using the built-in *icbm2tal* function (Lancaster et al., 2007). Statistical analysis of the transformed foci was validated using Monte Carlo Simulation (1,000 permutations) with cluster-forming voxel-level threshold at uncorrected *P* < 0.001 combined with cluster-size correction using family-wise error at *P < 0.05* (Eickhoff et al., 2016). Cluster statistics for overlapping ALE maps were generated with GingerALE using the Contrast Studies function. The algorithm assigns a value to each voxel that overlaps between two maps by reporting the minimum ALE value in the voxel between the two overlapping maps. The final *P* maps generated by GingerALE have been uploaded to NeuroVault, accessible at https://identifiers.org/neurovault.collection:5539 (Gorgolewski et al., 2015).

For any given clusters across each map, we report the following two coordinates: 1) weighted center (WC) and 2) extrema value (EV). Whereas the WC represents the centroid of a cluster, EVs represent a local maximum ALE score within a significant cluster. Anatomical labels of each coordinate were queried via the SPM Anatomy toolbox (Eickhoff et al., 2007, 2006, 2005). Finally, resulting ALE maps were displayed using multi-slice and surface views generated using Mango software (http://ric.uthscsa.edu/mango/) and the high-resolution MNI-space Colin 27 template provided by GingerALE (http://brainmap.org/ale/).

## 3. Results

### 3.1 Rhythm

The rhythm ALE analysis revealed significant clusters in both hemispheres of the brain, most of which were symmetrically mirrored (**Figure 2**, **Table 3**). The most notable clusters appeared bilaterally in the dorsolateral (pars Opercularis) part of the inferior frontal gyrus (IFG). Another pair of notable clusters emerged within the bilateral basal ganglia, including tissue from the caudate head, putamen, and the globus pallidus. Additional bilateral clusters emerged in the supplementary motor area (SMA)— including the pre-SMA—superior temporal gyrus (STG)/temporparietal junction (TPJ), inferior parietal lobule (IPL), and insula. Some areas emerged in a single hemisphere. For example, the premotor cortex (PMC) and precentral gyrus emerged in the right hemisphere, while the cerebellum (Crus I) appeared in the left hemisphere.

**Figure 2:**
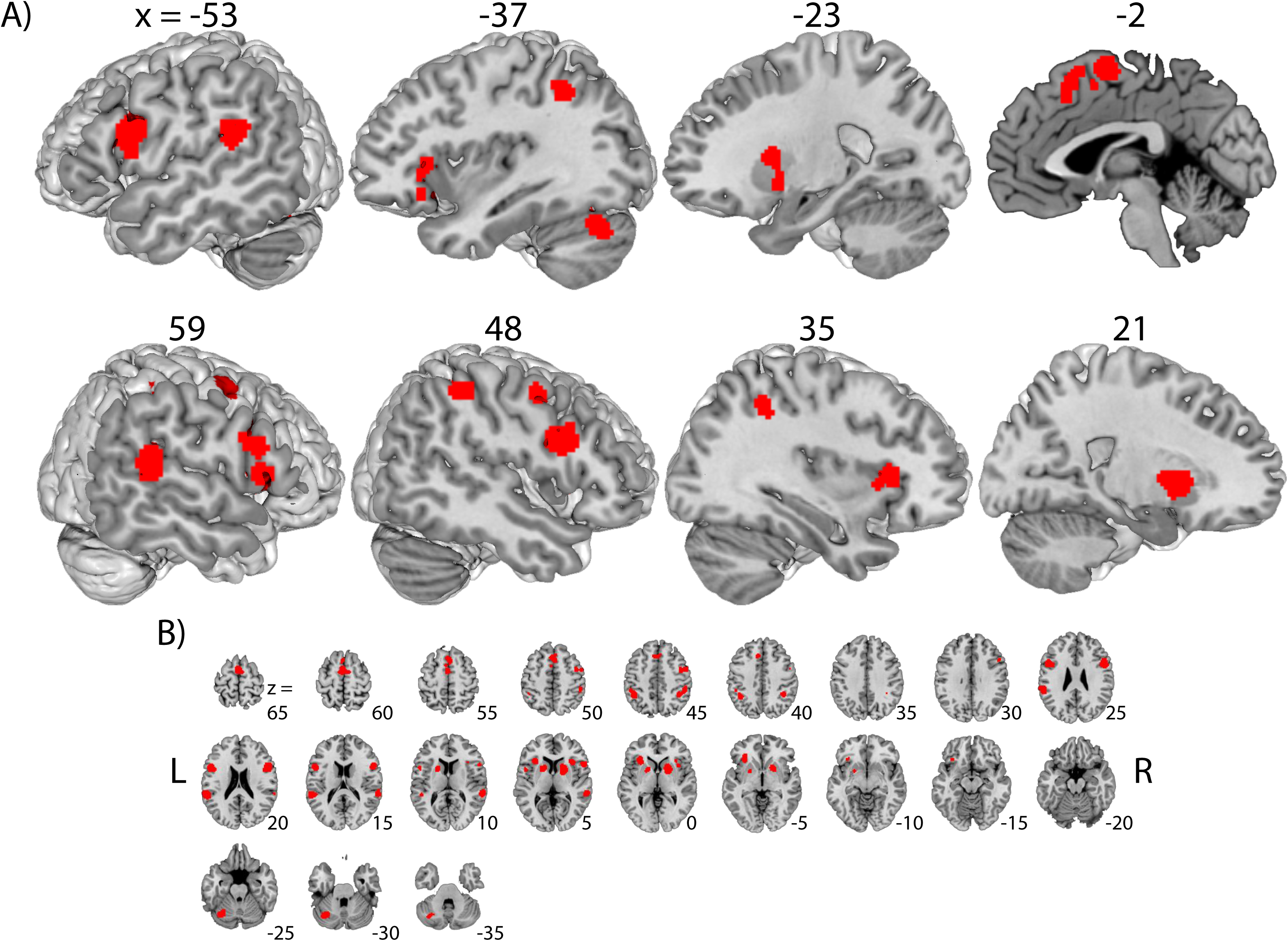
Results of rhythm analysis. (A) Rendering view of the rhythm ALE map at different slices. (B) An axial view of select slices to better illustrate rhythm clusters in cortical, sub-cortical, and cerebellar regions. The map was thresholded at voxel-level *P < 0.001* (uncorrected) in combination with cluster-level *P < 0.05* corrected using *FWE*.

**Table 3:**
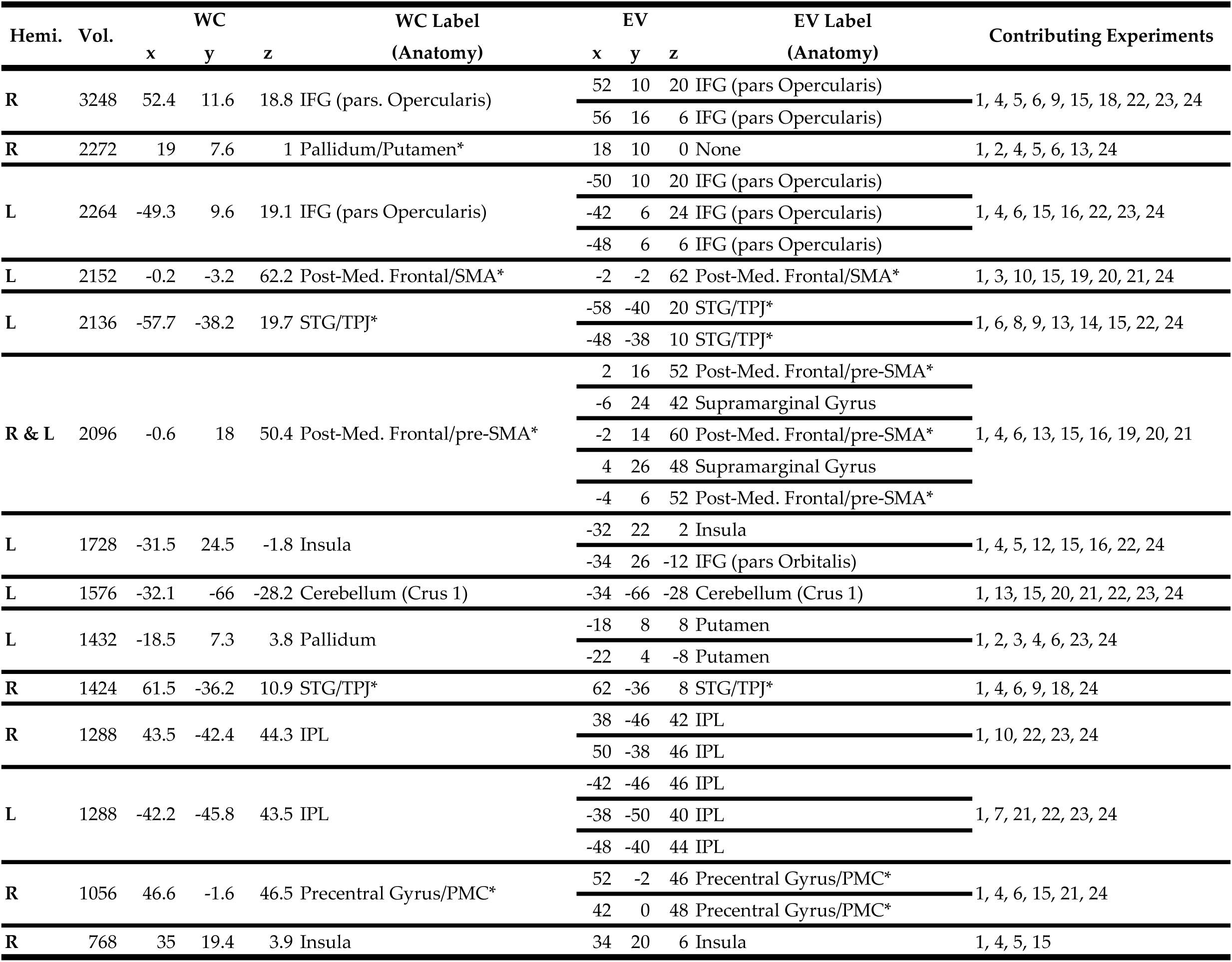
Rhythm clusters found by ALE analysis. WC: the weighted center of the cluster; EV: the extrema value; IFG: inferior frontal gyrus; Post-Med. Frontal: posterior-medial frontal; STG: superior temporal gyrus; TPJ: tempoparietal junction; IPL: intraparietal lobule; PMC: premotor cortex. Asterisk indicates that the second anatomical label was assigned manually by the authors.

### 3.2 Syntax

Unlike the rhythm ALE analysis, the syntax analysis yielded significant clusters predominantly in the left hemisphere with the exception of the right insula. The most notable cluster spanned both the pars Triangularis and Opercularis regions of the left IFG, running dorsally into the middle frontal gyrus and ventrally into the insula. Another notable cluster appeared in the middle and posterior aspects of the middle and superior temporal gyri. Outside of the fronto-temporal regions, sizable clusters were located in the SMA, left PMC, and left intraparietal lobule (IPL), yet no significant clusters emerged within the basal ganglia (**Figure 3A**, **Table 4**).

**Figure 3:**
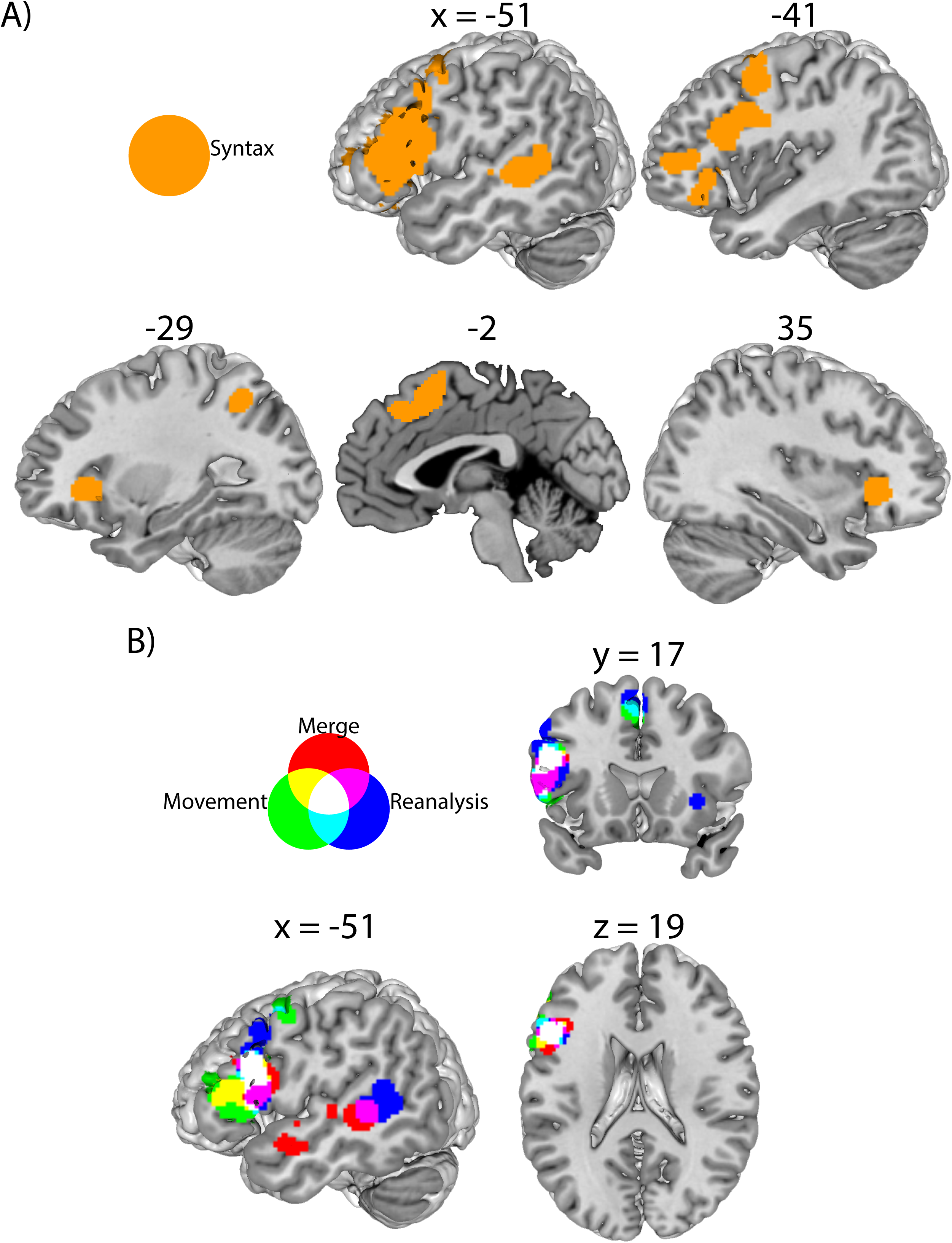
Areas engaged in syntax processes. (A) Renders of the combined syntax analysis, from the left and right hemisphere with slices (from left to right, top to bottom) x = [−51, −41, −29, −2, 35]. (B) Syntactic sub-types juxtaposed on the same rendering template. All maps were thresholded with voxel-level *P < 0.001* (uncorrected) in combination with cluster-level *P < 0.05* corrected using *FWE*. Data in (A) was generated from a down-sampled selection of syntax papers (see Methods), while (B) features all merge, movement, and reanalysis experiments.

**Table 4:**
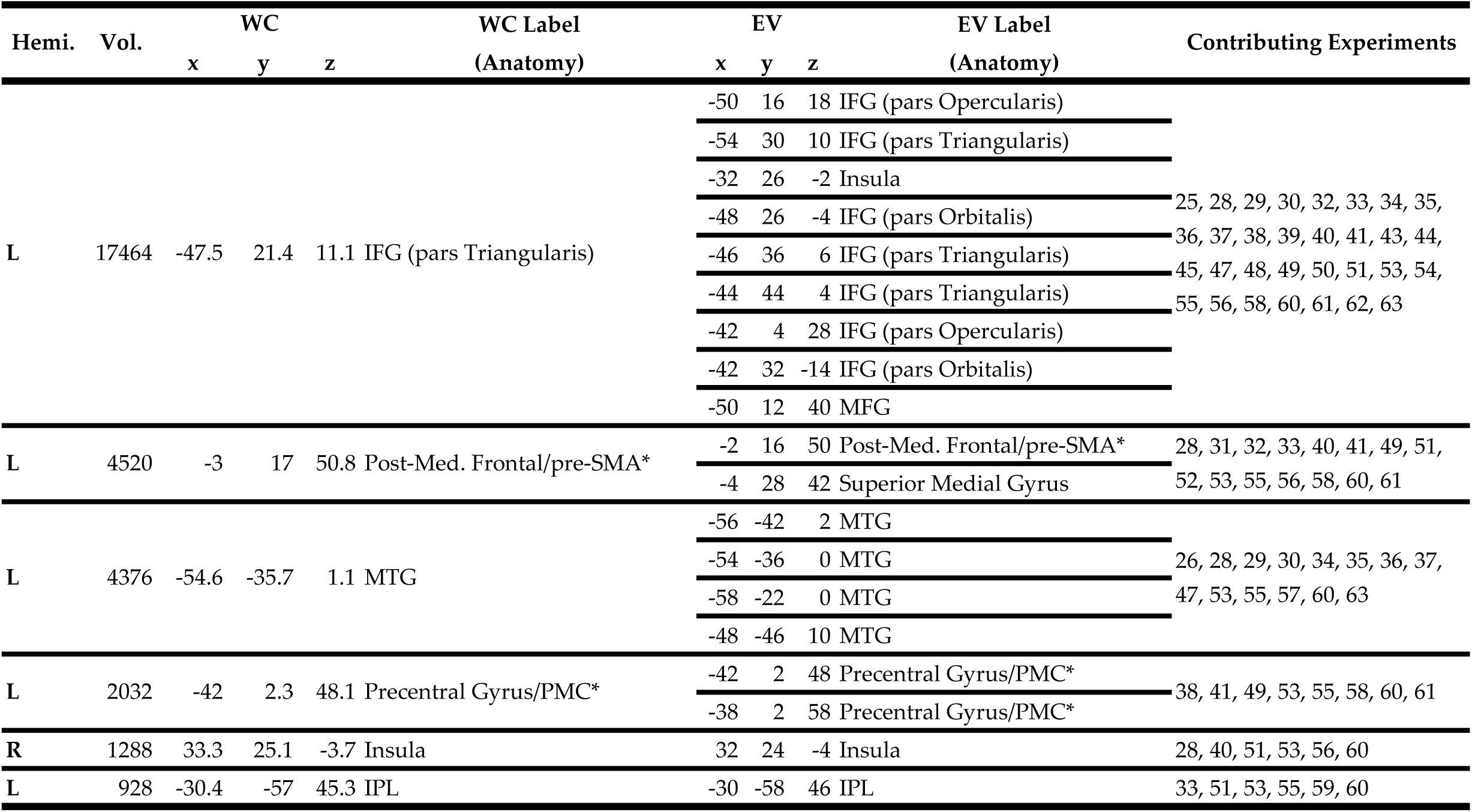
Clusters found by the combined syntax ALE analysis. WC: the weighted center of the cluster; EV: the extrema value; IFG: inferior frontal gyrus; MFG: middle frontal gyrus; MTG: middle temporal gyrus; STG: superior temporal gyrus; Post.-Med. Frontal: posterior-medial frontal; SMA: supplementary motor area; PMC: premotor cortex; IPL; intraparietal lobule. Asterisk indicates that the second anatomical label was assigned manually by the authors.

Next, we performed a series of ALE analyses on each subset of syntax experiments: merge, movement, and reanalysis (**Figure 3B**, **Table 4**). For merge, three significant clusters emerged within the left hemisphere: within the pars Triangularis of the IFG, the anterior STG, and posterior STG/MTG. For movement, significant clusters emerged in the left IFG encompassing both the pars Triangularis and Opercularis, the left PMC, and pre-SMA. Compared to the other two syntactic processes, reanalysis recruited more widespread regions. The largest cluster was found within the left IFG, which extended into the middle frontal gyrus and PMC. Several other regions were recruited including the posterior STG/MTG, pre-SMA, IPL, and PMC on the left hemisphere, as well as the right insula. Individual maps of merge, movement, and reanalysis can be found in the Supplementary Materials S3, S4, and S5.

These three sub-syntax maps were then overlaid to determine the common regions across different syntactic processes (**Figure 3B**). Tripartite overlap was seen mostly in the dorsolateral part of the left IFG (pars Opercularis, **Table 5**). However, there were several pair-wise overlaps throughout the left frontotemporal network. For example, in the left IFG, overlap was seen between merge and movement within the ventrolateral part (pars Triangularis). In SMA and PMC, overlap was seen between merge and reanalysis. In the temporal lobe, both merge and reanalysis recruit the posterior STG (**Table 5**).

**Table 5:**
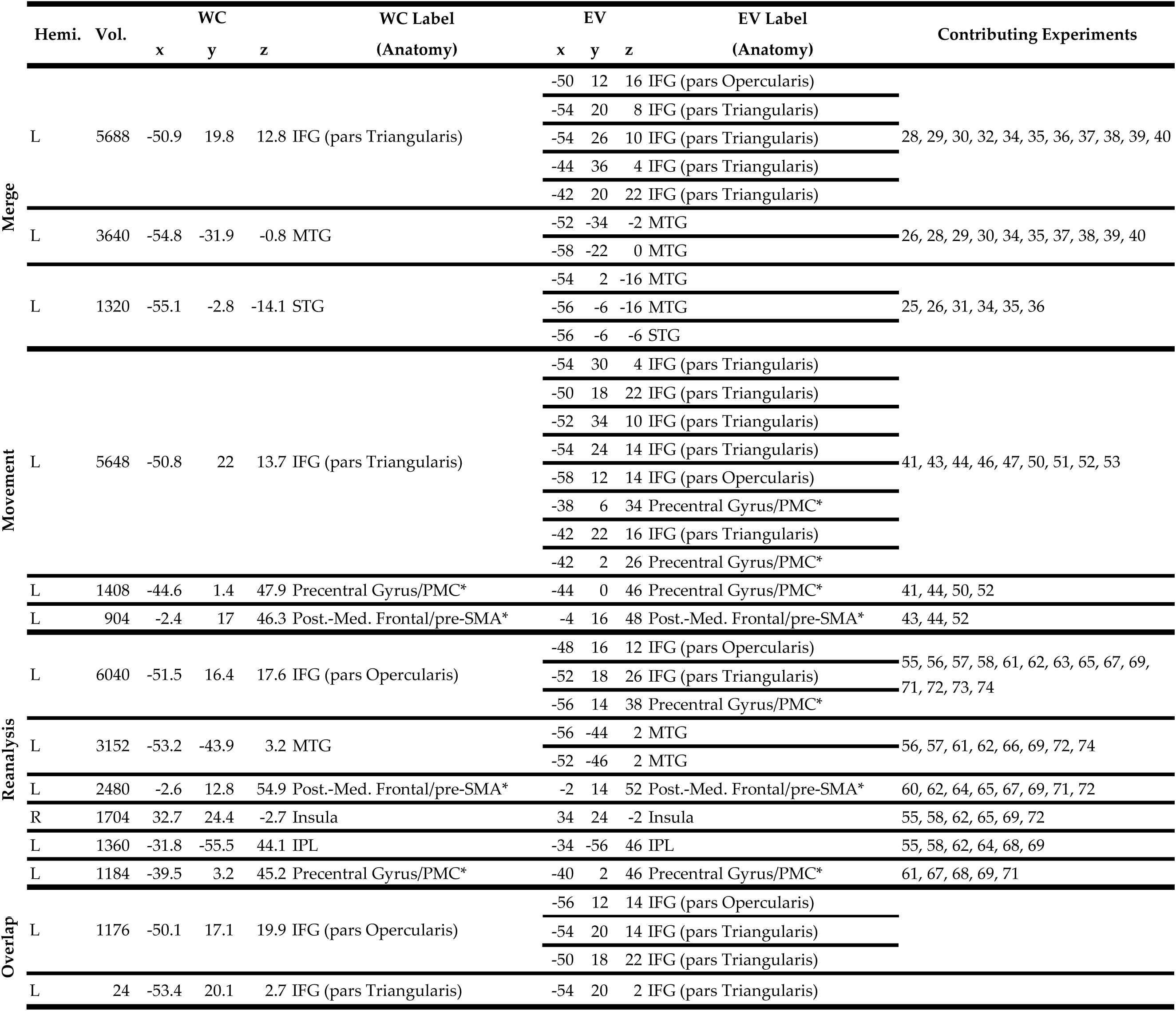
Syntax clusters found by ALE analysis of merge, movement, and reanalysis. WC: the weighted center of the cluster; EV: the extrema value; IFG: inferior frontal gyrus; MTG: middle temporal gyrus; STG: superior temporal gyrus; PMC: premotor cortex; Post.-Med. Frontal: posterior-medial frontal; SMA: supplementary motor area; IPL; intraparietal lobule. Asterisk indicates that the second anatomical label was assigned manually by the authors.

### 3.3 Overlap between Rhythm and Syntax

After we delineated ALE maps for both rhythm and syntax, we generated maps for overlap between the two. This revealed overlapping clusters in the dorsolateral part of the left IFG (pars Opercularis), the left SMA, and the bilateral insula with more pronounced activity in the left hemisphere. Although no overlap was seen in the posterior aspect of the left STG, a rhythm cluster was located just superior to a syntax cluster in this region (**Figure 4**, **Table 6**).

**Figure 4:**
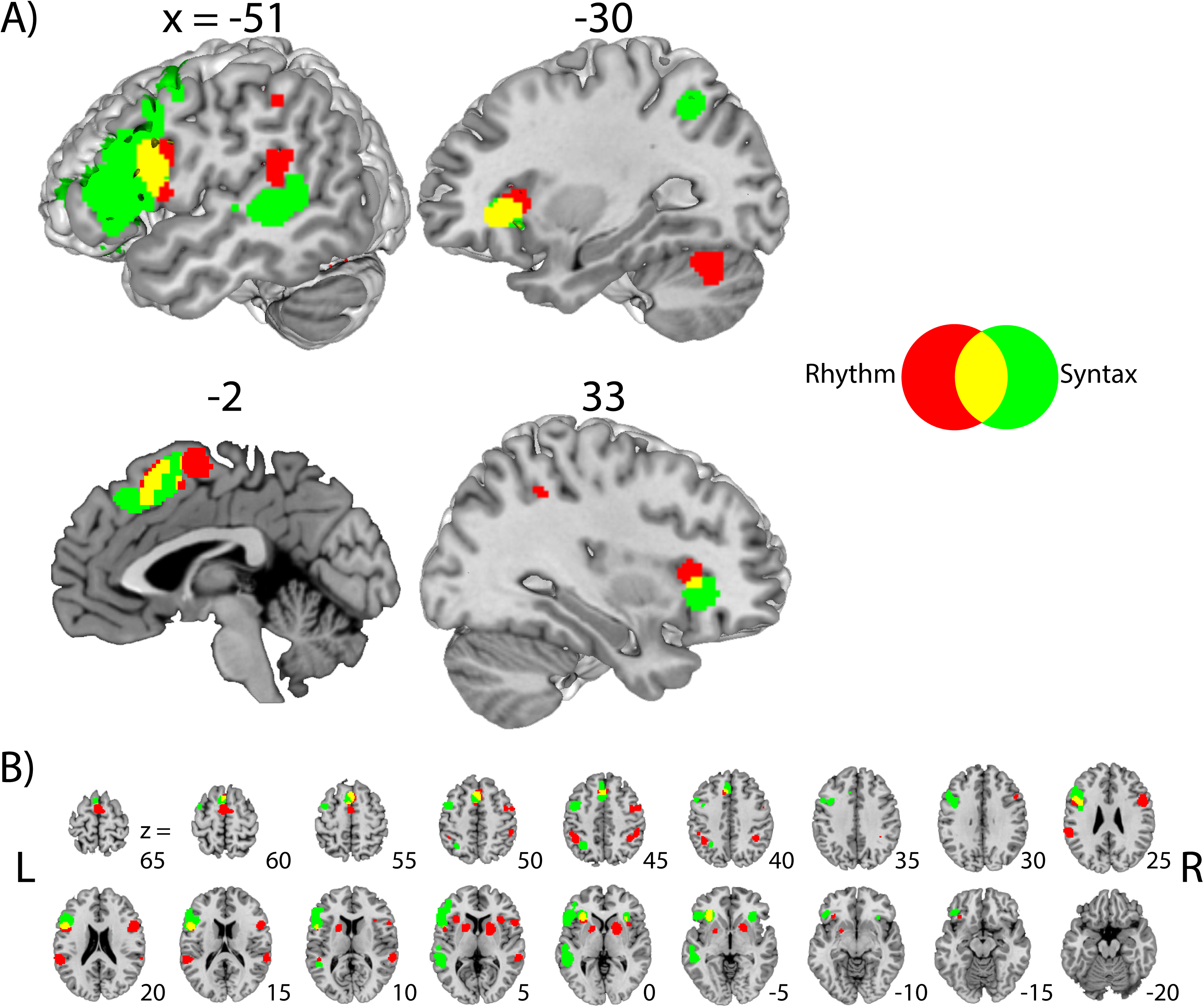
Overlap between rhythm and syntax analyses. (A) Renders of rhythm and syntax ALE maps at slices (from left to right, top to bottom) x = [−51, −30, −2, 33]. (B) A series of axial slices for rhythm and syntax areas. Both the rhythm and syntax maps were thresholded at voxel-level *P < 0.001* (uncorrected) in combination with cluster-level *P < 0.05* corrected using *FWE*.

**Table 6:**
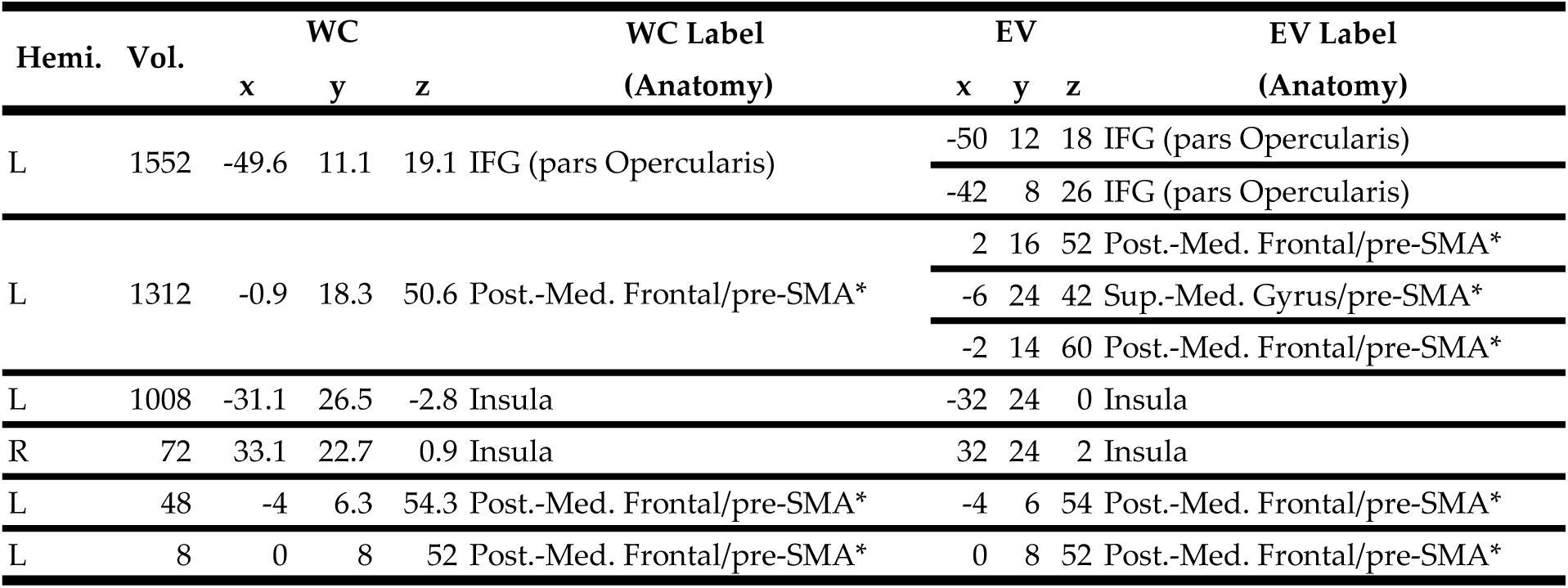
Overlapping clusters between rhythm and all syntax. WC: the weighted center of the cluster; EV: the extrema value; IFG: inferior frontal gyrus; Post.-Med. Frontal: posterior-medial frontal; Sup.-Med. Gyrus: superior medial gyrus; SMA: supplementary motor area. Asterisk indicates that the second anatomical label was assigned manually by the authors.

Next, we explored overlaps between rhythm and each syntax operation: merge, movement, and reanalysis (**Table 7**). Whereas merge and rhythm only shared a single cluster in the dorsolateral aspect of the left IFG (pars Opercularis, **Figure 5A**), both movement and reanalysis exhibited overlaps with rhythm in the left IFG (pars Opercularis) and the SMA (**Figure 5B**). Reanalysis also exhibited additional clusters in the IPL and right insula (**Figure 5C**).

**Figure 5:**
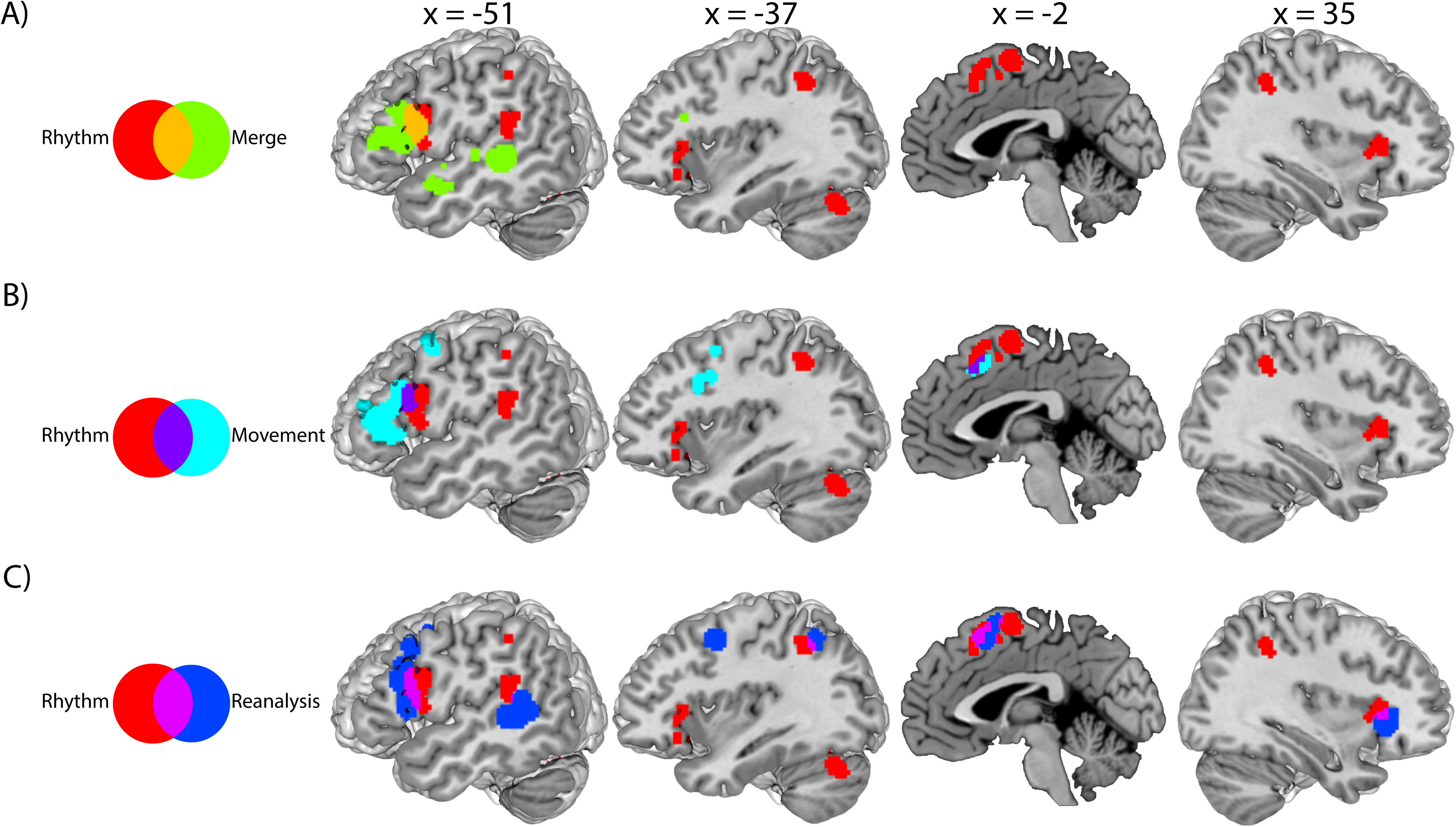
A series of sagittal slices depicting the degree of overlap between rhythm and each subset of syntax regions. (A) Rhythm and merge overlap at the pars Opercularis of the left IFG. (B) Rhythm and movement overlap in both the left IFG and pre-SMA. (C) Rhythm and reanalysis overlaps in the left IFG, left IPL, left SMA, and the right insula. All maps were thresholded at voxel-level *P < 0.001* (uncorrected) in combination with cluster-level *P < 0.05* corrected using *FWE*.

**Table 7:**
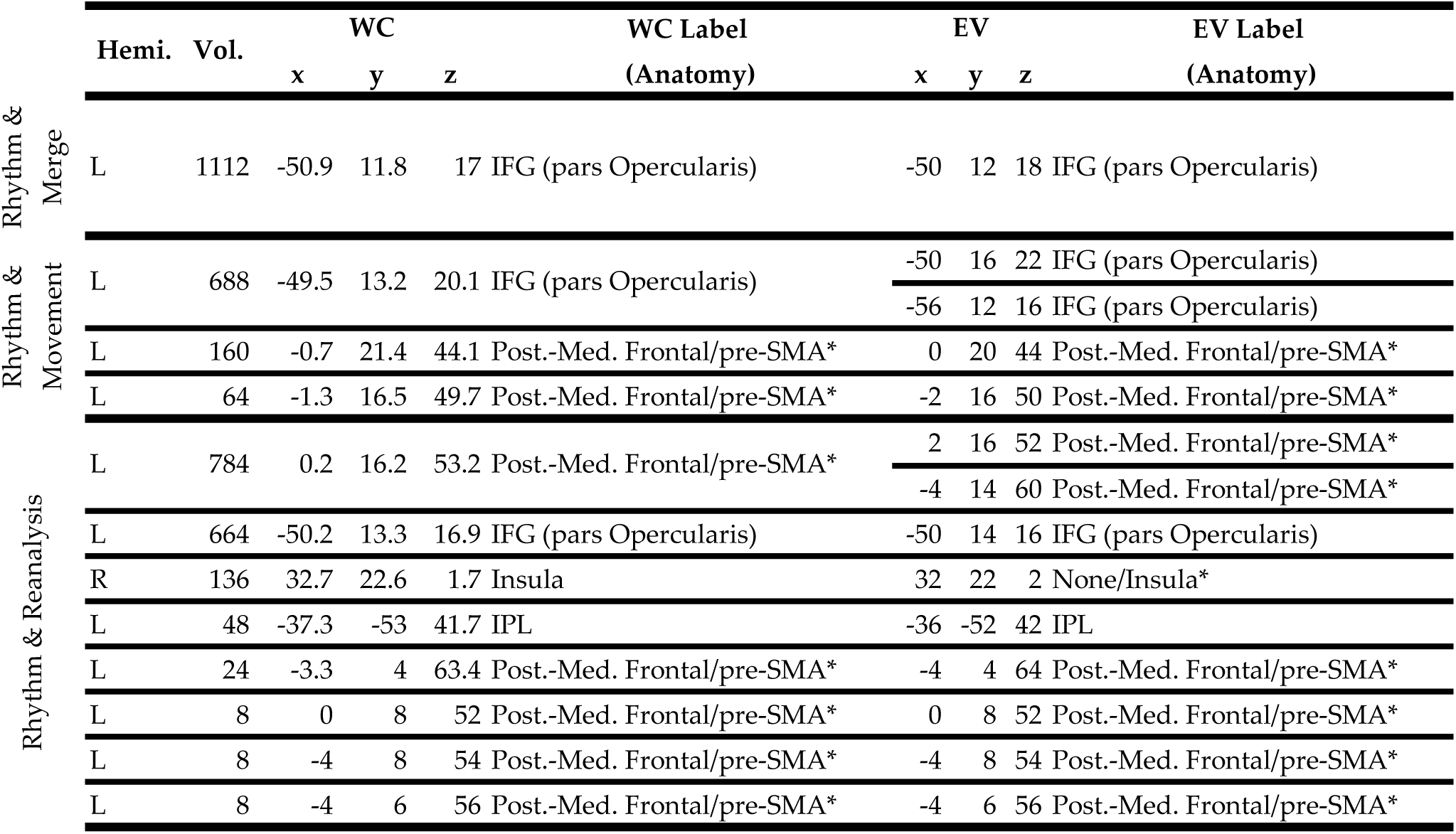
Overlapping clusters between rhythm and each syntax analysis. WC: the weighted center of the cluster; EV: the extrema value; IFG: inferior frontal gyrus; Post.-Med. Frontal: posterior-medial frontal; SMA: supplementary motor area; IPL: intraparietal lobule. Asterisk indicates that the second anatomical label was assigned manually by the authors.

## 4. Discussion

The primary goal of the present article is to provide a comprehensive overview of the brain-wide network that is recruited by both musical rhythm and linguistic syntax. To this end, we exhaustively searched for neuroimaging studies in these two domains. Out of an initial 2,504 neuroimaging studies considered by the present analysis, we found 24 rhythm experiments pertaining to rhythm, beat, and meter as well as 50 syntax experiments pertaining to merge, movement, and reanalysis that qualified for a series of ALE meta-analyses. Rhythm mostly recruited symmetrical clusters in bilateral cortical and subcortical areas including the IFG, putamen, SMA, STG, insula, and IPL. By contrast, syntax predominantly engaged a left-lateralized network including the IFG, PMC, STG, insula, and IPL. Overlap between rhythm and syntax was found in the left IFG, left supplementary motor area, and bilateral insula. Additional intersections between rhythm and each syntax process yielded clusters within a similar part of the left IFG (pars Opercularis), but only movement and reanalysis recruited motor regions such as the SMA. In what follows, we discuss how the current findings can shed light on the theoretical framework and behavioral evidence suggesting connections between music and language.

### 4.1 Overlap between Rhythm and Syntax

There is a growing body of behavioral evidence indicating a substantial influence of rhythmic and timing context on syntactic processing. For example, in Jung et al (2015, 2016), participants read garden path sentences that were presented word-by-word in an isochronous fashion while listening to chord sequences. When the critical disambiguating word was presented off-beat compared to the rest of the sentence, participants had a more difficult time parsing sentences (Jung, 2016; Jung et al., 2015). Conversely, it has been shown with various children populations that coherent rhythmic and metric cues facilitate identification of morphosyntactically correct and incorrect sentences. These populations include children with speech and language deficits such as dyslexia (Przybylski et al., 2013; Schmidt-Kassow and Kotz, 2008b, 2008a), SLI (Bedoin et al., 2016), and cochlear implants (Bedoin et al., 2017), as well as typically-developing children (Bedoin et al., 2017, 2016; Przybylski et al., 2013). Moreover, priming with rhythmic sequences containing easy-to-extract meter facilitated language comprehension compared to priming with rhythms that induced a complex meter (Przybylski et al., 2013). This facilitation effect does not appear to transfer to math and visuospatial tasks (Chern et al., 2018). Beyond these examples of interference and facilitation, Bedoin and colleagues (2017) demonstrated a transfer of training between the two domains by developing a rhythm training program to restore syntax comprehension abilities for congenitally deaf children with cochlear implants.

These behavioral connections between rhythm and syntax have been supported by neuroimaging data, mostly from electroencephalography (EEG; Kotz and Gunter, 2015; Roncaglia-Denissen et al., 2013). For example, German listeners understood syntactically ambiguous sentences more accurately when sentences were spoken with a regular versus irregular meter (Roncaglia-Denissen et al., 2013). The behavioral facilitation was accompanied by reduction of the P600 component—a hallmark of syntactic reanalysis (Frisch et al., 2002)—suggesting an interaction between meter and syntax processing. Relatedly, a case study using EEG with a Parkinsonian patient showed that the P600 component was restored priming with a regular beat (i.e. marching music; Kotz and Gunter, 2015). Although the exact source of this P600 remains elusive, the present study suggests that the left IFG, SMA, and bilateral insula may be candidate loci whose activities can be modulated by external beat priming during syntactic processing. That is, these areas may incorporate rhythmic (temporal) context during moment-to-moment analysis of syntactic structures. Such neural sensitivity to timing may be sharpened through music training, which may confer benefits to language (Patel, 2014) by enhancing temporal processing and/or predictive coding in these regions.

Alternatively, could the observed functional overlaps between rhythm and syntax reflect generic cognitive processes that are recruited by any type of task involving attention, decision-making, and working memory? Indeed, most of these overlapping clusters are parts of the so-called “extrinsic mode network (EMN)” that are active in a non-specific task-generic manner (Hugdahl et al., 2015). However, these frontal loci have been implicated in numerous neuroimaging studies of music and language; there is a firm consensus that these regions are integral parts of these respective networks (Friederici, 2018; Kunert et al., 2015; Nguyen et al., 2018). Because the EMN includes fronto-parietal-temporal cortices, it is difficult to completely rule out a possibility that some of the overlapping clusters reported here may have appeared due to generic cognitive processes that both rhythm and syntax entail. For this reason, we have taken our due diligence to guard against reports of non-specific ALE activation. For example, we included experiments contrasting rhythmic and syntactic tasks versus an active baseline (e.g. meter tapping versus beat tapping, syntactically difficult versus easy sentences) instead of resting baselines. Furthermore, we applied a stringent statistical threshold (*P < 0.001* voxel-wise in combination with *P < 0.05* family-wise error cluster size correction) in the resulting rhythm and syntax map.

Nevertheless, we acknowledge that some of the overlapping frontal clusters may reflect non-specific cognitive resources recruited by both domains.

Conversely, the fact that the same voxels are activated by rhythm and syntax does not necessarily indicate that the same neurons are involved in the two tasks, as there are hundreds of thousands of neurons and glia within a voxel (Koelsch, 2011). This is an intrinsically challenging problem that current state-of-the-art neuroimaging research still faces. Within the overlapping cluster, it is plausible that some neurons are exclusively responding to musical rhythm or linguistic syntax processing, but not both. Future studies with higher resolution neuroimaging as well as focal stimulation using advanced transcranial magnetic stimulation) or electrocorticography will allow researchers to better identify at a finer-grained scale the neuronal populations that are tuned to the common or distinct aspects of rhythm and syntax.

### 4.2 Temporal Hierarchy Processing

An important characteristic of music and language is the hierarchical organization of serial temporal information (Fadiga et al., 2009; Fiebach and Schubotz, 2006; Fitch and Martins, 2014; Jackendoff, 2009; Jeon, 2014; Lashley, 1951). For example, along the hierarchy of rhythm structure, the lowest unit consists of the onsets of tones (i.e. rhythm), from which a pulse or beat is extracted, which in turn gives rise to meter (Fitch, 2013; Lerdahl and Jackendoff, 1983; Vuust and Witek, 2014; Zatorre and Zarate, 2012). Similarly, along the hierarchy of grammar, word roots can join affixes to form new roots (e.g. establish; establish*ment*; *dis*establish*ment*), giving rise to differences in word meanings (Jackendoff, 2009). A set of words then form syntactic phrases that ultimately are the constituents of a whole sentence. Many studies examining temporal hierarchy in syntax have implicated the left IFG in both the music and language domains (for a review, see Fitch and Martins, 2014). For example, the left IFG was shown to be responsive to violations of linguistic and musical syntax (Fedorenko et al., 2009; Koelsch et al., 2005b; Slevc et al., 2009; Sun et al., 2018). In the present study, we found significant overlap between musical rhythm and linguistic syntax, which is in line with these previous findings.

Importantly, temporal hierarchies can also account for action sequencing in the motor domain (Fadiga et al., 2009; Fiebach and Schubotz, 2006; Fitch and Martins, 2014; Jackendoff, 2009; Jeon, 2014; Lashley, 1951; Pulvermüller, 2014); according to Jackendoff’s model (Jackendoff, 2007), complex actions can be broken down into three components: preparation, head, and coda. The head forms the core of an action and contains steps to execute the main goal. Preparation, however, consists of any steps that need to be completed before execution of the head can start. Lastly, codas are any series of steps required to return the system to normalcy. Such action sequences are mainly mediated by frontal motor circuitries including the left IFG, PMC, and SMA (Clerget et al., 2009; Koechlin and Jubault, 2006). These frontal motoric processors exhibited overlaps between rhythm and syntax in the present study, suggesting that they are also at play in analyzing the temporal hierarchies for musical rhythm and linguistic syntax. Relatedly, early left anterior negativities (ELAN) and parietal P600 components were observed when canonical structures were violated in action and language (Maffongelli et al., 2015). More recently, Casado et al. (2018) reported interaction of both ELAN and P600 between syntax and complex motor processes. Although the overlapping clusters were found mostly within the left hemisphere in the present ALE study, some studies have implicated either the bilateral or the right frontal cortex for music (Cheung et al., 2018; Farbood et al., 2015; Koelsch et al., 2013, 2005a; Maess et al., 2001), language (Bahlmann et al., 2008), and action (Koechlin and Jubault, 2006; Schubotz and Cramon, 2004). For example, a recent fMRI study created stimuli that violated temporal structure at multiple levels of organization by scrambling a famous piano concerto. While a variety of auditory areas were activated in response to the scrambled concerto, the right IFG was solely active when professional pianists listened to the unscrambled piece (Farbood et al., 2015). Together, the frontal motor network appears to be involved in the dynamics of temporal structure pervasive in music, language, and action.

### 4.3 Predictive Coding

Both rhythm and syntax entail predictive coding in order to more efficiently process upcoming events (Koelsch et al., 2019; Kuperberg and Jaeger, 2016; Rohrmeier and Koelsch, 2012; Staub, 2015; van der Steen et al., 2013; Vuust and Witek, 2014). For example, in music, listeners have a tendency to tap before the actual beat (Repp, 2005; Repp and Su, 2013), indicating that listeners are actively predicting the next beat. Indeed, the sensation of syncopation is achieved by disrupting active predictions (Vuust and Witek, 2014), a phenomenon which is often used by composers on purpose. Linguistic syntax also leverages prediction during comprehension. Syntactic surprisal paradigms are particularly based upon estimating how likely a next word fits the canonical syntactic structure (Hale, 2001; Levy, 2008). While such paradigms have been used in previous neuroimaging studies, there were not enough studies that qualified to be included in the present ALE analysis. However, predictive coding can be considered in the data obtained from syntactic merge, movement, and reanalysis. For merge, it is more likely for a determinant to be followed by a noun (e.g. {the} + {car}) than it is to be followed by a word that typically functions as a verb (e.g. {the} + {jump}). For movement, sentences beginning with ‘Wh-’ should give rise to prediction. Lastly, as an example of reanalysis, garden pathing occurs when predictions are violated.

What neural substrates are responsible for predictive coding in rhythm and syntax? Past neuroimaging studies have highlighted the putamen, SMA (Grahn and Rowe, 2013) and PMC (Jantzen et al., 2007) in tasks involving prediction for rhythm sequences. Similarly, words with high syntactic surprisal cause a response in the right putamen, bilateral IFG, and insula (Henderson et al., 2016). Together, these results demonstrate how fronto-striatal networks play a crucial role in predicting structure in musical rhythm and linguistic syntax (Kotz et al., 2009). Such observations in music have led to the Action Simulation for Auditory Processing hypothesis (Iversen et al., 2009; Iversen and Balasubramaniam, 2016; Patel and Iversen, 2014) proposing that action processing is recruited during predictive coding of music and language.

### 4.4 Overlap in Merge, Movement, and Reanalysis

Although our primary goal was to identify overlapping clusters between musical rhythm and linguistic syntax using ALE data, we attempted to make additional comparisons between different types of syntactic processes by taking advantage of the large number of neuroimaging experiments on merge, movement, and reanalysis. Of course, there are many other types of syntactic operations, such as surprisal (Henderson et al., 2016), and morphosyntactic transformations (Sahin et al., 2006). Unfortunately, too few experiments in these domains qualified for the ALE analysis. However, this is an important avenue to be explored by future ALE studies when more neuroimaging studies of syntactic surprisal have accumulated.

The extra comparisons within syntax yielded a single tripartite cluster only in the pars Opercularis, one of the constituting parts of Broca’s area. Emerging evidence garnered from neuroimaging and neurophysiology studies have indicated functional (Sahin et al., 2009) and anatomical segregation between the pars Opercularis and pars Triangularis—the two sub-units of Broca’s area (Amunts et al., 2010). Importantly, these two adjacent areas have been hypothesized to handle different types of language operations; the pars Triangularis has been implicated in the “semantic combinatorics” (Friederici, 2018) required in sentence comprehension, as well as lexical decision tasks (Heim et al., 2005). The pars Opercularis has been implicated in general sequencing and hierarchical processing in linguistic syntax (Friederici, 2018, 2002), which may explain the overlap among the three syntactic processes observed in the current study.

Beyond the left IFG, movement and reanalysis engaged the PMC and SMA. While both regions have been implicated in auditory and language processing (Hertrich et al., 2016; Lima et al., 2016), their participation in syntax has received less attention compared to the IFG and STG. Our data revealed that involvement of the PMC and SMA depends on the type of syntactic process being studied. In order to resolve violations of prediction or to rapidly process grammatical rules, these motor areas may require more coordination with the left IFG and PMC/SMA via the aslant tract (Dick et al., 2014; Vassal et al., 2014). Future studies are warranted to elucidate the neuroanatomical connection between these frontal areas during various types of syntactic processing.

## 5. Conclusion

Over the past few decades, increasing evidence has been garnered regarding the behavioral connections between music and language in general, as well as between rhythm and syntax in particular (Gordon et al., 2015a). The present ALE meta-analysis attempted to lay the groundwork demonstrating detailed neuroanatomical overlap between musical rhythm and linguistic syntax. Our findings well speak to hierarchical processing (Fitch and Martins, 2014), temporal prediction (Vuust and Witek, 2014), and sequencing (Kotz et al., 2009); processes that are mediated by the frontal motor circuitries including left IFG, SMA, and bilateral insula.

## Acknowledgements

Thanks to Simon Eickhoff for answering methodological questions about ALE and GingerALE software. We thank members in the SLAM lab for their helpful comments on the earlier version of the manuscript. This study was funded by Center for Brain Injury, part of The Ohio State University Discovery Initiative and the neuroimaging pilot grant from the College of Arts and Sciences at OSU.

