## Supplementary figures and images for "Shared neural resources of rhythm and syntax: An ALE Meta-Analysis"

### Supplementary Figure 1

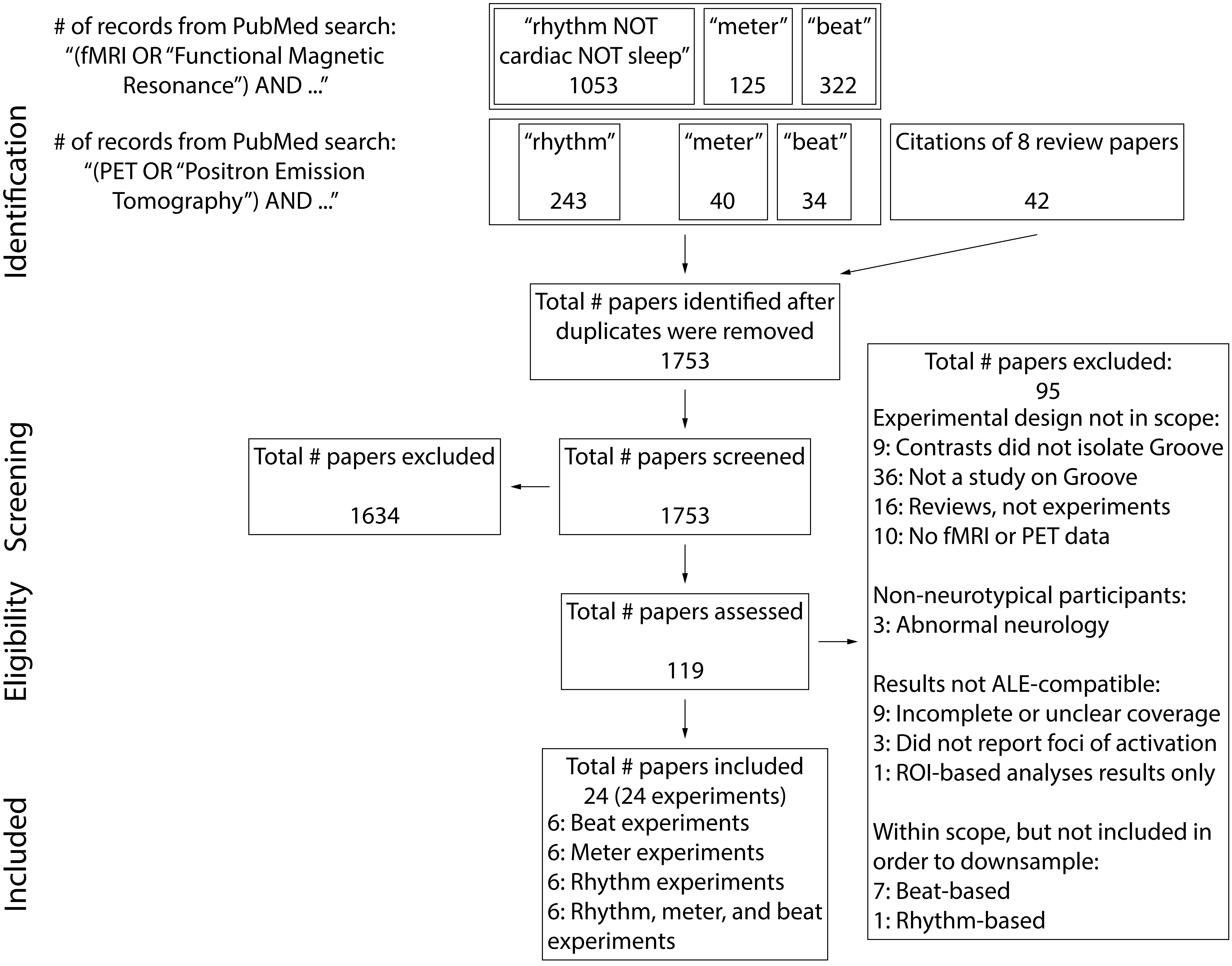

### Supplementary Figure 2

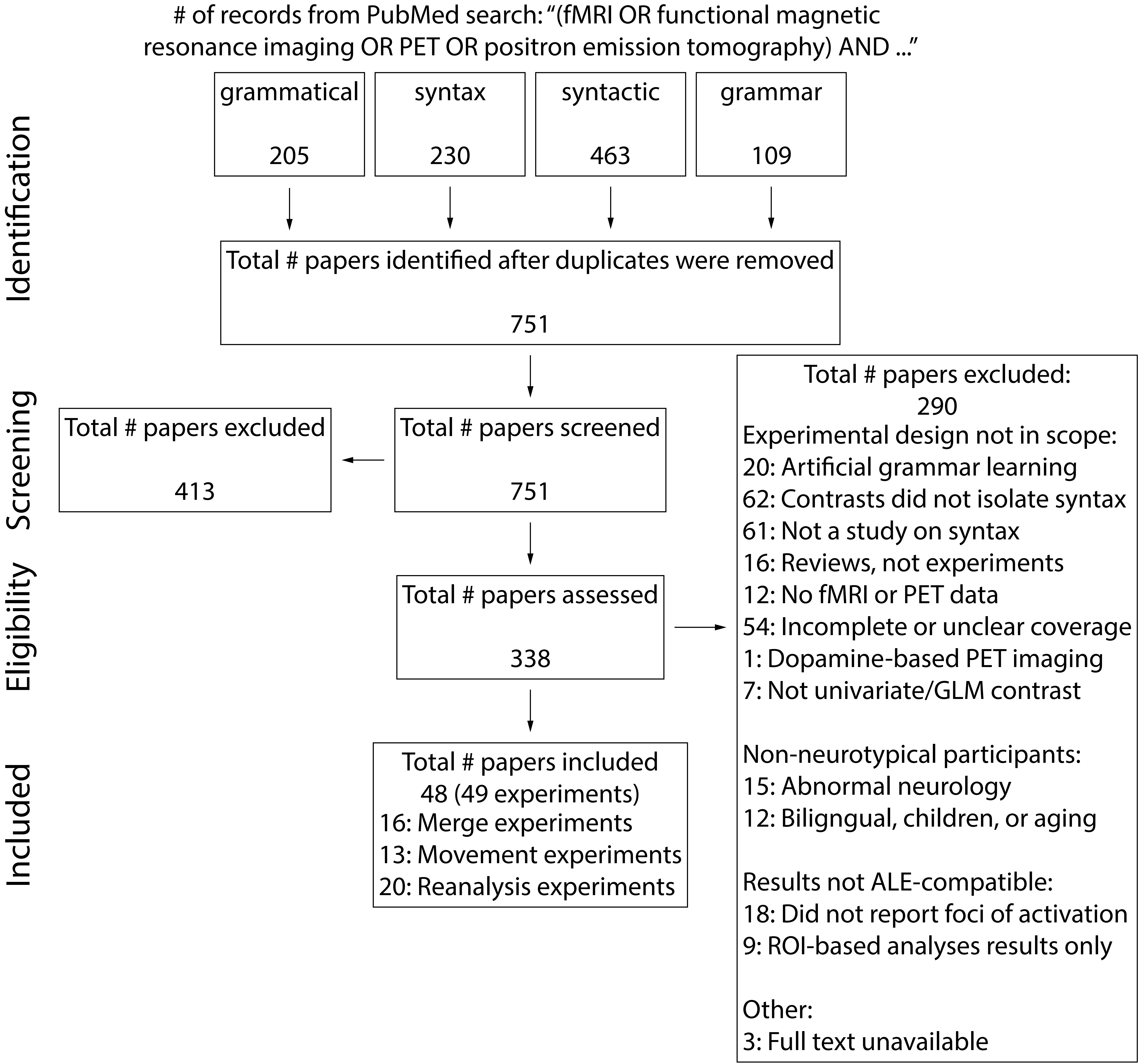

### Supplementary Figure 3

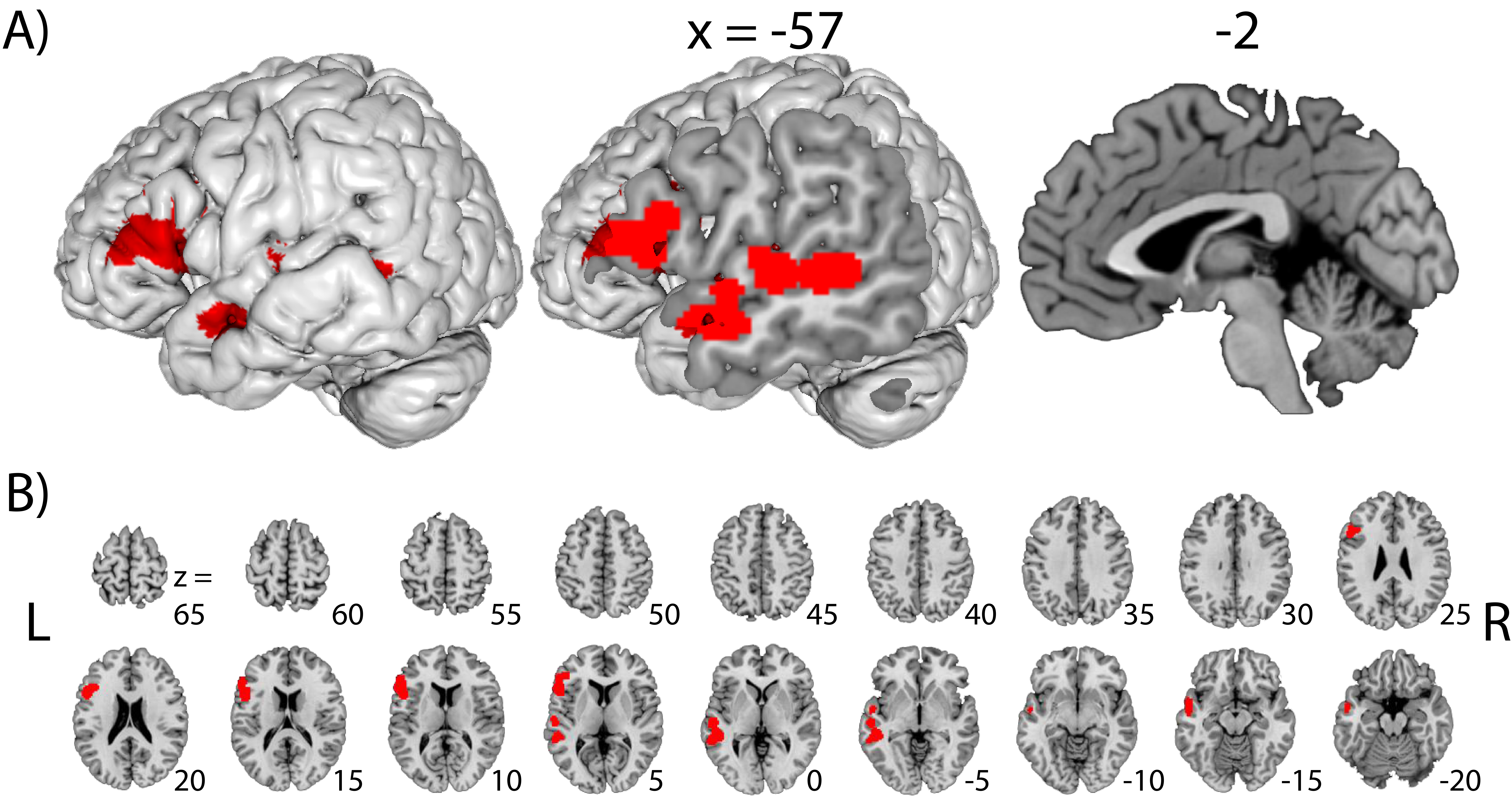

### Supplementary Figure 4

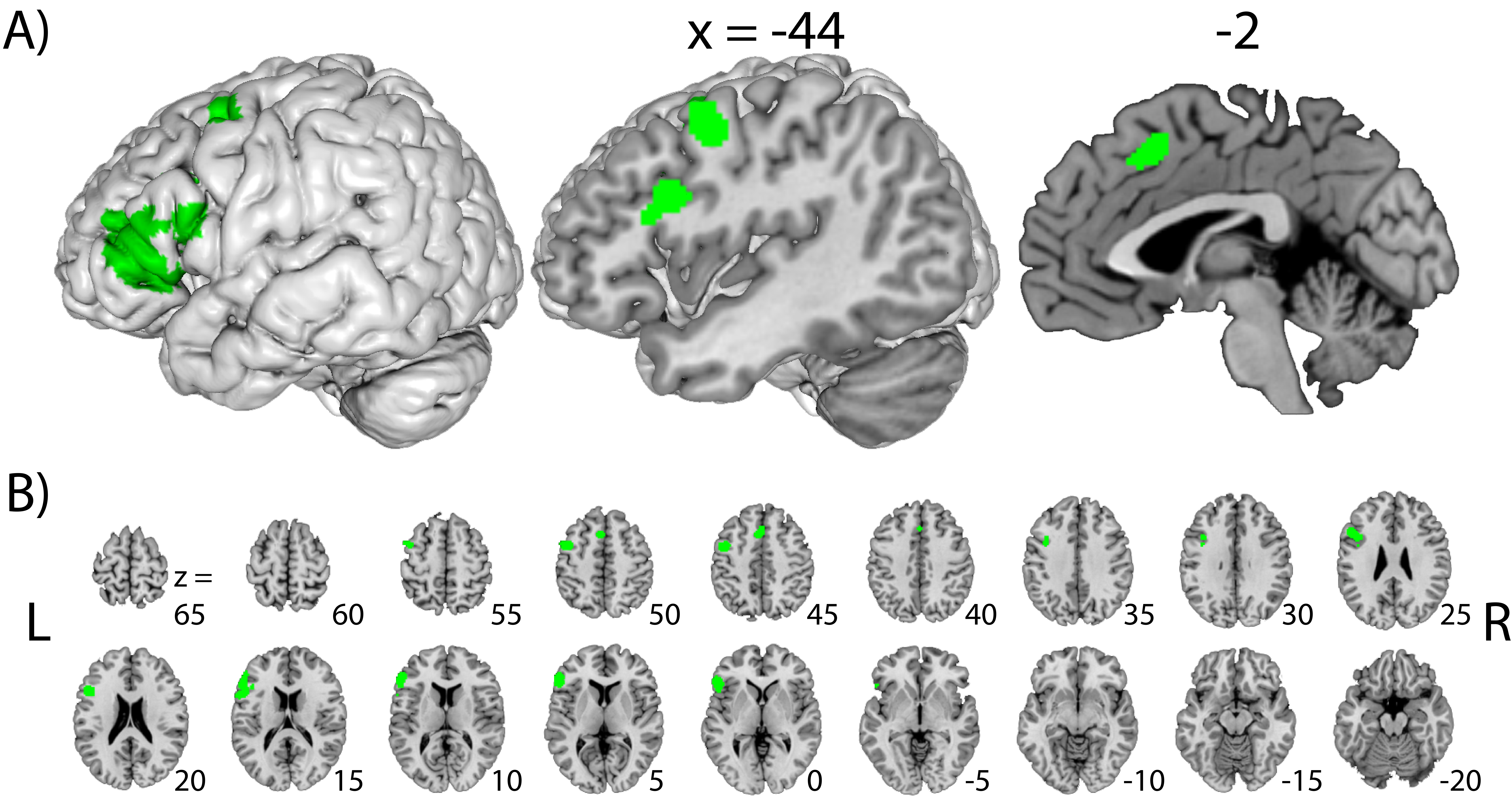

### Supplementary Figure 5

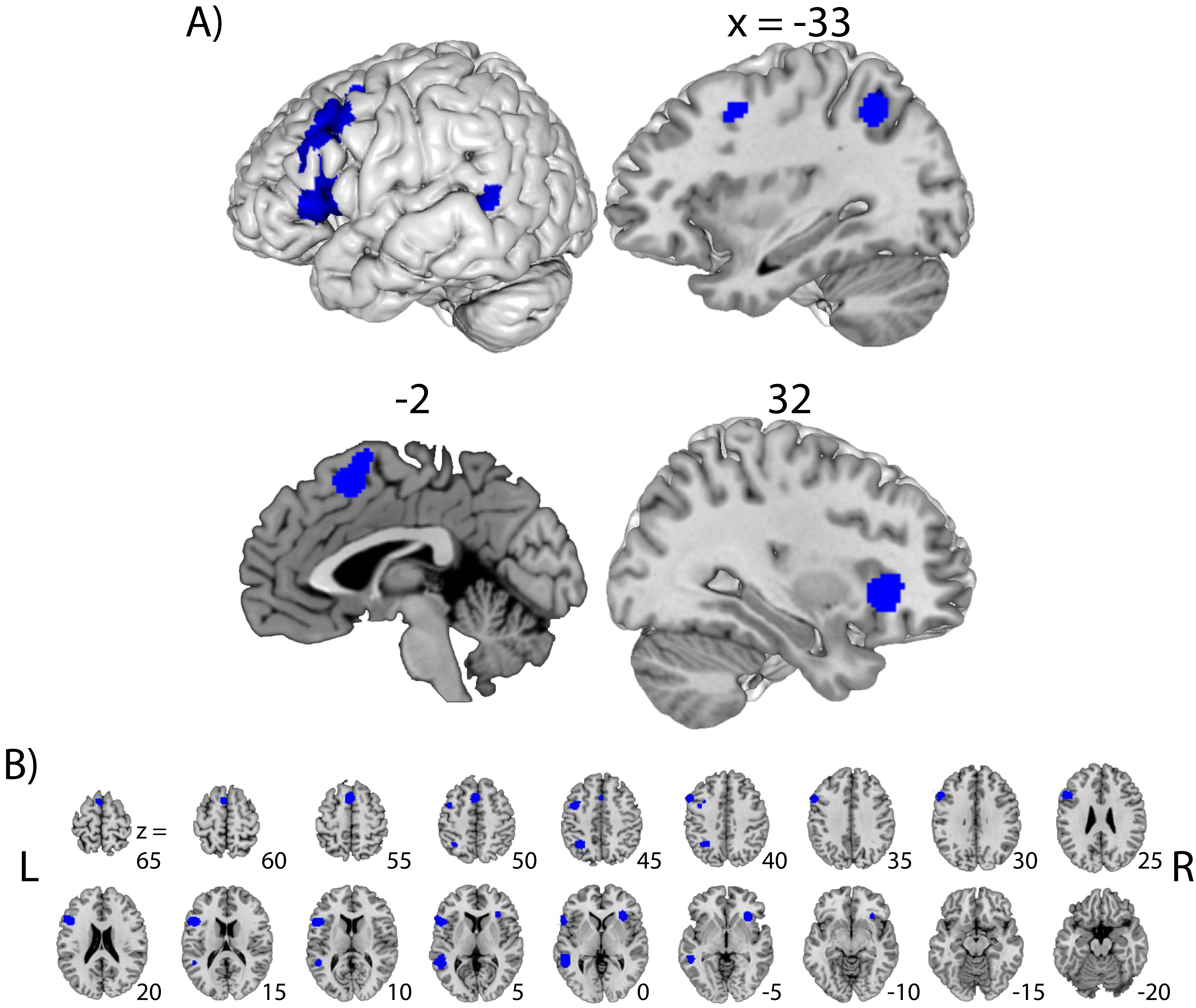
